# Novel chemolithotrophic and anoxygenic phototrophic genomes extracted from ice-covered boreal lakes

**DOI:** 10.1101/139212

**Authors:** Lucas Sinclair, Sari Peura, Pilar Hernández, Martha Schattenhofer, Alexander Eiler

## Abstract

Although an important fraction of the world’s lakes remains ice-covered during a large proportion of the year, little is known about the microorganisms that govern the biogeochemical processes occurring under-ice along the stratigraphic redox gradients. Reconstructed genomes provide evidence for anoxygenic photosynthesis involving fixation of carbon using reduced sulphur and iron as an electron donor in the anoxic zone of the sampled lake systems. In addition to anoxygenic photosynthesis, our molecular data reveals novel chemolithoautotrophic organisms and supports the existence of methanotrophs in bottom anoxic waters. Reconstructed genomes matched methanotrophs related to *Methylobacter* tundripaludum, phototrophic *Chloroflexi* and *Chlorobia*, as well as lithoautotrophic genomes affiliated to the *Betaproteobacteria* class and *Planctomycetes* phylum. Based on our in-depth characterization, complex metabolic interactomes emerge unique to each lake’s redox tower and with sulfur, iron and carbon cycling tightly intertwined through chemolithotrophy and anoxygenic photosynthesis.

## Introduction

Lakes around the globe have a pronounced impact on the global carbon cycle [1] [2]. For example, the estimated carbon losses through outgassing and burial from inland waters reach the same magnitude as total global net ecosystem production [2]. Many of the processes related to carbon cycling rest to a large extent on poorly understood microbial processes. Green-house gas (GHG) emissions from seasonally ice-covered water systems at high latitude have been argued to substantially contribute to global GHG production [3]. Moreover, these systems are thought to be particularly sensitive to climate change as future reductions in the duration of ice-cover are estimated to increase annual water body emissions by 20–54% before the end of the century [4]. Still, scientific attention remains primarily focused on their ice-free state [5] while lakes frozen temporarily are not put ‘on hold’ when ice-covered in the winter season. A recent study [6] found that, in twelve small lakes in subarctic Sweden, the *CO*_2_ emitted at ice-melt accounted for 12 to 56% of the annual *CO*_2_ emitted from these lakes. Understanding microbial processes under ice is thus essential to construct accurate models and predict the lake systems’ role in global biogeochemical cycles such as those concerning the production of escaping GHGs including *CO*_2_, *CH*_4_ and *N*_2_*O*.

Although the physical properties of ice-covered systems are well known and predictive models have been developed [7], how exactly this change of state affects the biogeochemistry and microbial processes is poorly described and understood. Indeed, few studies have looked at parameters such as diversity, growth rate or metabolic capabilities of microbes under ice [5]. Work so far suggests that under-ice conditions greatly influence the diversity of bacterial communities via shifts in the availability and quality of organic matter and nutrients [8] [9].

Once ice-covered, hydrodynamic processes switch abruptly away from of the open-water period as exchanges with the atmosphere are halted, including nutrient depositions and GHG emissions. Light input is reduced, particularly when a layer of snow forms on top of the ice. In addition to reduced mixing, the temperature gradient reverses in the opposite direction to that of summer, with surface water switching from warmest to coolest layer, while the temperature in the bottom layer is rather constant through the year [7]. Notably, the creation of redox-depth gradients is expected to select for microbes with specific traits adapted towards the available electron acceptors and donors [10].

Despite their global significance for GHG emissions, the microbial processes occurring in the oxygen depleted or even anoxic ice-covered water column are as yet unknown and opaque. To what extent nitrogen, sulfur and iron cycling or fermentation regulate the degradation of organic matter and how phototrophic and lithotrophic processes intertwine with organic matter degradation by changing the availability of electron acceptors, are matters of ongoing debate. Although numerous metabolic processes can possibly occur under ice, such as methanotrophy, photoferrotrophy or the coupling of denitrification and iron oxidation, the molecular and geochemical evidence has so far been slim, with few exceptions as in [9] [11].

The principal aims of this study were to retrieve the phylogenetic composition of the microbial communities proliferating under ice in high latitudinal lakes. Using a combination of geochemical measurements and metagenomic techniques, we set out to disentangle the biogeochemical processes along depth profiles of five Swedish lakes, with certain resemblance of the million boreal lakes in Fennoscandia, Siberia and the Canadian shield [12]. By reconstructing the genomes of the abundant microbes we revealed the genomic basis of metabolic traits previously undescribed in these poorly studied environments, such as photoferrotrophy and methanotrophy. Finally, through linking the metabolic potential of the microbes to the cycling of elements, we propose that sulfur, iron and carbon cycling are tightly linked through anoxygenic photosynthesis and chemolithotrophy in these systems.

## Results

### Correspondence of bacterial diversity with lake characteristics

All five sampled lakes were thermally stratified with four of the systems displaying steep redox gradients as indicated by oxygen, iron and sulfate profiles (figure 1, supplementary figure S2). At the water-ice interface, methane bubbles were observed in two of the systems, LB and BT, with *CH*_4_ concentrations above 5 μM. The profiles also indicated accumulation of methane in lake RL with a reading of 150 nM *CH*_4_. Nevertheless, methane concentrations were highest in the water column just above the sediment layer, reaching far beyond 50 μM in two systems (LB and KT), similar to the maximum concentrations previously reported for boreal lakes [13]. Carbon dioxide followed similar trends with concentrations increasing with depth and maximum values ranging from 0.22 to 0.95 mM in the five systems.

**Figure 1.**
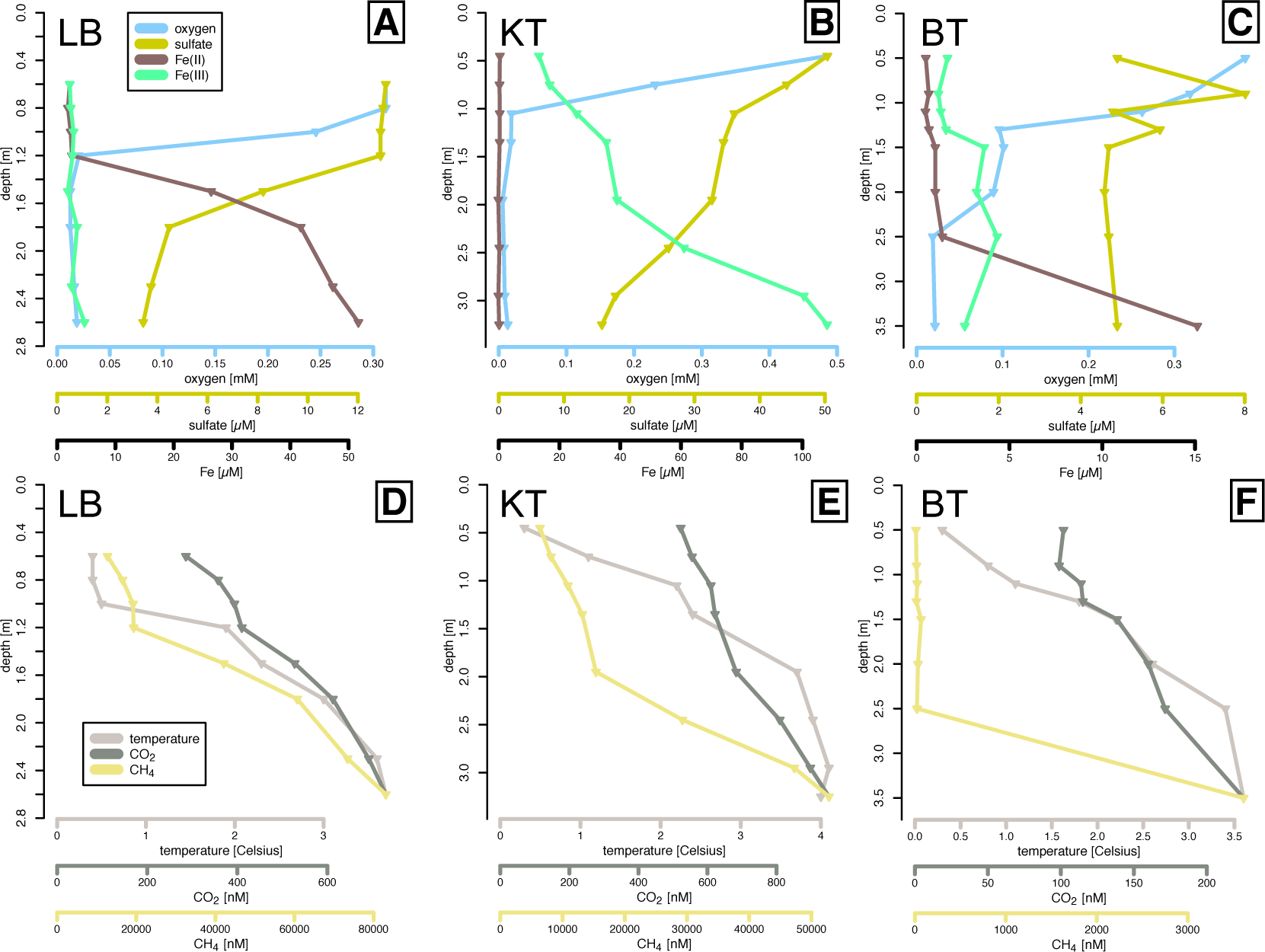
Summary of lake characteristics. Profiles of water chemistry (A-C) and gas concentrations (D-F) in lakes LB (A,C), KT (B,D) and BT (C,E). For RL and SB see supplementary figure S2.

Clear differences between the lakes concerning maximum concentrations and profiles of inorganic nutrients and electron acceptors such as oxygen, sulfate and Fe(III) were evident. Lake SB was the most oligotrophic system with very low total organic carbon (TOC) and nutrient concentrations. There was no oxygen gradient, but the water column was oxygenated throughout. Surface oxygen concentration close to saturation (13.8 mg/l at 2 °C) in RL and SB suggest active oxygenic photosynthesis in these systems. Lakes BT and RL had oxygen concentration below detection limit only in the deepest 1 m, while in LB and KT, which represent the most productive systems as indicated by the bacterial cell numbers (supplementary figure S1), oxygen was already depleted at 1 m depth from the surface. In the latter two, sulfate concentrations decreased with depth, suggesting the presence of sulfate-based anaerobic respiration. Whereas iron occurred in its reduced form (FeII), it occurred exclusively in its oxidized form (FeIII) in KT with concentrations above 100 μM even in oxygen-depleted layers.

These differences in chemical profiles among the studied systems were also reflected in the unique bacterial communities present in individual lakes as assessed by 16S rRNA gene amplicon sequencing. As visualized in the ordination plot in figure 2A, depth profiles of individual lakes grouped within lake with no convergence at any depths. This selection of microbial communities by the specific conditions of each system is further emphasized by the observation that environmental parameters significantly relate to community patterns. Indeed, pH, Fe(II), Fe(III) and total organic nitrogen (TON) together explained 47% of the observed variability as inferred by a redundancy analysis (p < 0.001). Further evidence is thus provided for the tight link between stratigraphic patterns in taxonomic composition and the depth-related variations in redox potential.

**Figure 2.**
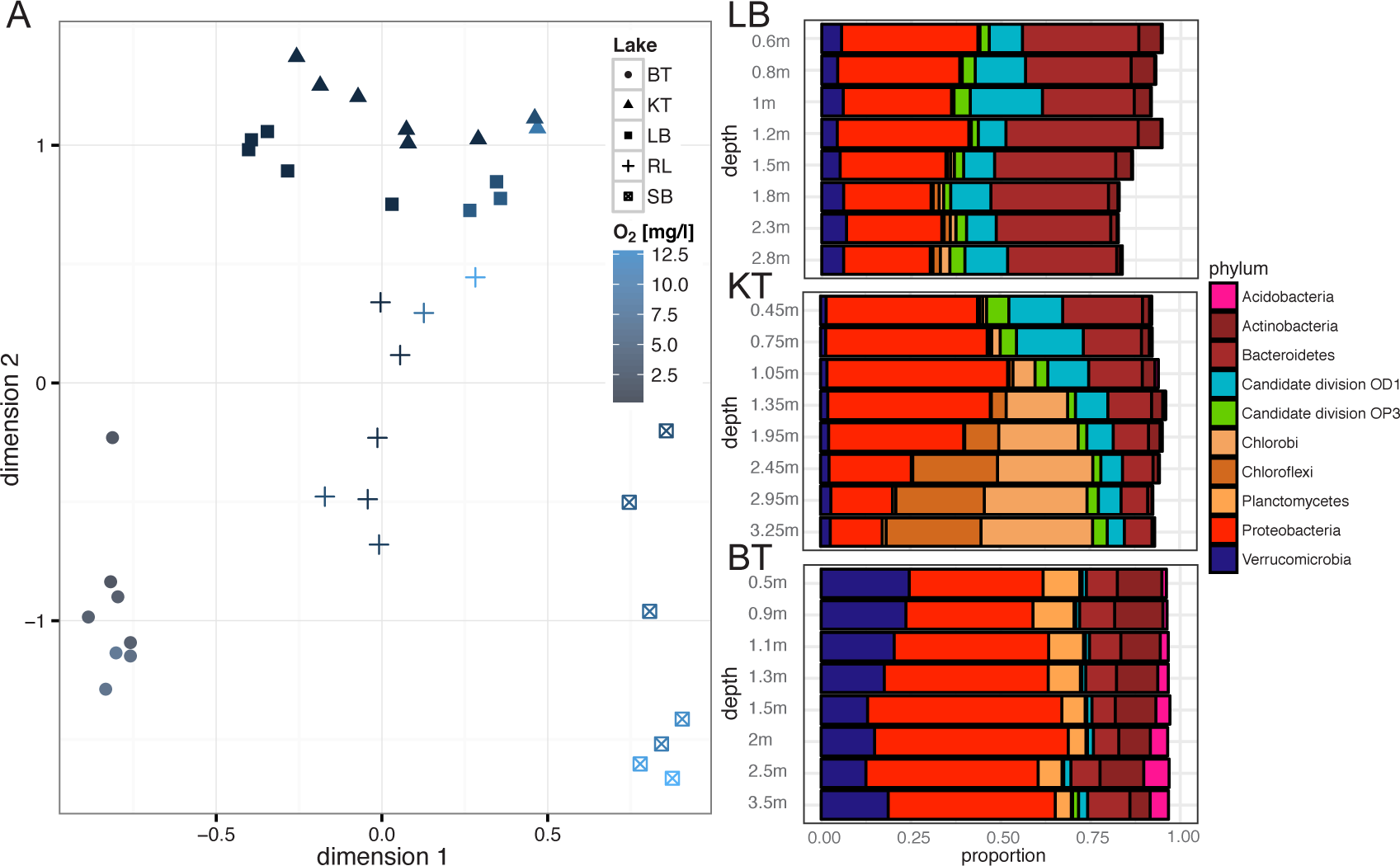
Bacterial community profiles by lake and by depth. Non-parametric multidimensional scaling plot (A) and depth profiles of individual lakes LB, KT and BT with the ten most abundant phyla. For RL and SB, see supplementary figure S3.

Overall, 44 phyla of which fifteen candidate phyla were identified across all samples. Proteobacteria, *Actinobacteria* and *Bacteroidetes* dominated in the five systems with differences among the lakes and distinct stratigraphic patterns as seen in figure 2. The fifteen candidate phyla recruited on average 12.8% of the reads ranging from 2.6 to 28.5% per sample and these percentages increased toward the lake bottoms, as previously observed [14]. The most prominent vertical redox-gradient related changes were the increase in the relative abundance of *Chlorobia* and *Chloroflexi* in KT, and a shift towards *Deltaproteobacteria*, Candidate division BSV13 and *Lentisphaera* at intermediate depths in LB. Specific taxonomic groups identified were indicative for aerobic (e.g. family *Comamonadaceae*) and anaerobic anoxygenic photosynthesis (e.g. phylum *Chlorobia*, and genus *Oscillochloridaceae*), recalcitrant polymer degradation (e.g. genera *Paludibacter* and *Chthoniobacter*) and methanotrophy (e.g. families *Methylococcales*, *Methylophilaceae* and genus *Candidatus Methylacidiphilum*) as seen in figure 2 and in supplementary figure S3. This detailed taxonomic analysis showed partial concordance with previously described boreal lake communities (for comparison see [14] [15] [16] [8])

### Microbial traits along three under ice redox gradients

We used the more detailed trait predictions offered by the shotgun-metagenomics technique to infer the functional potential of a multitude of uncultured freshwater prokaryotes and gain insight into their potential role in elemental cycles. Using the chemical information gathered (in particular iron and sulfate concentrations) as well as the bacterial community depth profiles built (in particular the proportions of candidate phyla and other poorly described deep branching taxonomic groups), we chose three systems: LB, KT and BT. We obtained shotgun metagenome libraries resulting in a total of almost 530 million reads ranging from 15.1 to 26.5 million reads per sample. Assembly of individual depth profiles (i.e. individual lakes) resulted in almost 90’000 contigs with a minimum length of 1 kb, as seen in table 1. These contigs recruited 30.6% of the total reads with a range from 22.3 to 44.7% in individual systems. Thus, on average, 70% of the lake’s genomic content and the functions associated are not captured, introducing uncertainties in our metabolic trait profiles. More details on the different stages of the sequence processing are offered in online supplementary reports.

**Table 1.** Summary of sampling sites and sequencing results.

|  | Latitude<br>N | Longitude<br>E | TOC<br>[mg/l] | TOP<br>[µg/l] | Amplicon<br>reads | Shotgun<br>reads | Contigs |
| --- | --- | --- | --- | --- | --- | --- | --- |
| **BT** | 63.5821 | 12.2708 | 7.8 - 10.1 | 9.5 - 14.6 | 685'057 | 179'710'871 | 44'165 |
| **RL** | 63.5843 | 12.2743 | 7.8 - 7.0 | 5.4 - 14.1 | 811'181 | n/a | n/a |
| **LB** | 63.3382 | 12.5480 | 5.9 - 12.5 | 8.4 - 9.2 | 782'317 | 184'786'957 | 21'392 |
| **KT** | 63.3493 | 14.4587 | 15.8 - 26.6 | 12.7 - 111 | 842'872 | 165'342'647 | 23'668 |
| **SB** | 63.3961 | 13.1619 | 4.6 - 3.9 | 2.7 - 3.0 | 964'981 | n/a | n/a |

The final assemblies and mappings were used in combination with hidden Markov models (HMM) similarity searches to provide more information on the functional repertoires along the depth gradients. By screening the assembled metagenomes for genes encoding conserved protein family domains (PFAMs) of relevant enzymes (see supplementary table S2), we aimed to determine the metabolic potential of the key microbial processes in these boreal lakes. Although these annotations are derived from incomplete databases built on results from environments in which the majority of taxa have not been well characterized [17], we were able to note that the predicted functional diversity contained within the under-ice microbial communities was congruent with the taxonomic diversity. We show a strong positive correlation between the functional diversity, as assessed by PFAM annotations, and the taxonomic diversity derived by amplicon sequencing of the 16S rRNA gene in each of the three systems’ depth gradients. Indeed, the correlation coefficients were 0.70 in LB, 0.78 in BT and 0.77 in KT with p < 0.001 as determined by a procrustes superimposition. Although there has been considerable debate in the field of ecology as to how taxonomic diversity of communities relates to functional or trait-level diversity [18], recent studies on microbial communities from soils [19] and aquatic systems [20] have provided evidence for such a coupling, despite high functional redundancy in microbial communities [21].

Reflecting the unique taxonomic profiles of the individual lakes, the three selected systems were clearly separated when it came to their functional profiles as seen in supplementary figure S4. Still, there were common features along the depth profiles as revealed by the progression of genes encoding for methanotrophy, iron and sulfur oxidation-reduction reactions, phenolic compound degradation, anoxygenic photosynthesis and methanogenesis as seen in figure 3. The genes encoding sulfur and iron cycling were notably very abundant and genome equivalence indicated potential for sulfur and iron oxidation or reduction in over 50% of the microorganisms in all samples. The presence of sulfur oxidation genes (*sox* and *dsr*), together with steep gradients in sulfate concentrations and identified taxa such as *Geobacter*, *Desulfobulbus* and *Desulfovibrio*, as well as *Chlorobium*, point to an important role of sulfur cycling in the water column of these freshwater systems. Despite concentrations being much lower than those of marine systems, the importance of sulfur cycling has already been suggested for lake sediments [22]. Still, LB and BT were shown to have low sulfate and high iron concentrations as do many lakes in the boreal landscape.

**Figure 3.**
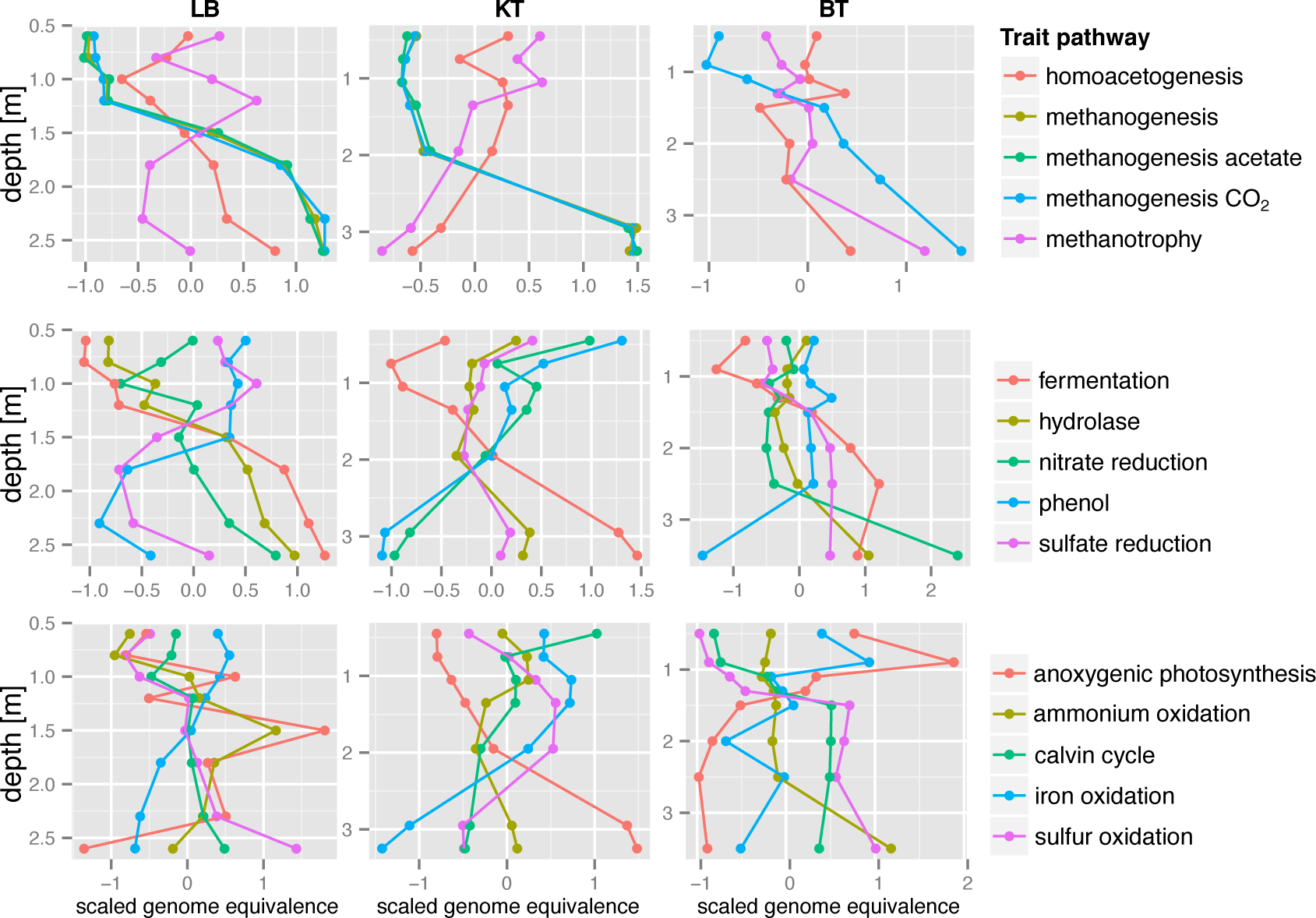
Genome encoded metabolic profiles by depth. Depth profiles visualizing the abundances of PFAM markers related to carbon and energy metabolism. The average of all scaled (z-score) HMMhits are indicative for particularmetabolic traits were plotted along with depth.

Annotations also revealed genes implicated in the reduction of sulfate/sulfite (*dsr*) and Fe(III) (*fer*), as well as denitrification genes (*nar*, *nir*, *nor* and *nos*) as part of chemolithotrophic redox reactions or anaerobic respiration of organic matter. Screening for known genes containing PFAM domains that catalyze the degradation of allochthonous organic matter revealed that the potential to hydrolyze plant polymers (e.g. cellulose and hemicellulose) could be found throughout the water column while organisms capable of degrading phenolic compounds (e.g. lignin) were mainly found in the oxic portion of the water column.

Genes involved in several fermentative pathways were detected with a genome equivalence value of approximately 1, indicating more than 1 copy per genome, in the deepest strata of LB and KT, suggesting a high potential for the production of fermentation products in bottom waters of boreal lakes. Likewise, genes encoding the terminal hydrogenase of *H*_2_-evolving fermentations (hydA) increased with depth. The abundance of formyltetrahydrofolate synthetase genes (fhs), which encodes the key enzyme of the Acetyl-CoA pathway of homoacetogenesis, was highest just below the ice and the oxycline as seen in figure 3 showing scaled absolute genome equivalent values close to 0.5 [23]. Key genes indicative for methanogenesis increased with depth together with the fraction of methanogenic *Archaea*. Three different orders of methanogenic archaea were found, *Methanobacteriales*, *Methanomicrobiales* and *Methanosarcinales*. The first two are hydrogenotrophic, producing *CH*_4_ from *H*_2_ and *CO*_2_, whereas *Methanosarcinales* is metabolically more versatile carrying out hydrogenotrophic, acetoclastic and methylotrophic methanogenesis. The presence of these different types of methanogens was also verified by the homology to key enzymes described in these processes including glutathione-independent formaldehyde dehydrogenase (FdhA), hydrogenase subunit A (EchA), formylmethanofuran dehydrogenase subunit A (FmdA), formylmethanofuran-tetrahydromethanopterin N-formyltransferase (FTR), methenyltetrahydromethanopterin cyclohydrolase (MCH), methylenete-trahydromethanopterin dehydrogenase (MTD), coenzyme F420-dependent N5, N10-methenyltetrahydromethanopterin reductase (MCH), tetrahydromethanopterin S-methyltransferase (MtrA), methyl-Co(III) methanol-specific corrinoid protein coenzyme M methyltransferase (MtaA), methyl-coenzyme M reductase alpha subunit (McrA), acetate kinase (AckA), acetyl-CoA synthetase (ACCS), phosphate acetyltransferase (PTA), heterodisulfide reductase subunit A (HdrA), acetyl-CoA decarbonylase/synthase complex subunit beta (CdhC). While co-occurring in all three systems, alternation from acetoclastic to hydrogenotrophic methanogens along the depth profiles differed among the three systems. Differences in the distribution of acetotrophic, methylotrophic and hydrogenotrophic methanogens pointed to a variability in the concentrations of fermentation products. Such differences can be indicative of variable efficiency in the fermentation steps, possibly owing to established interactions between fermentative syntrophic bacteria and their methanogenic counterparts.

The genes indicative for methanotrophy were detected throughout the watercolumn of all three lakes as seen in figure 3. Type I methanotrophs (RuMP based) of the families *Methylococcaceae* and *Methylophilaceae* were abundant in all three lakes in bottom samples (maximum of 14.4 and 20.9%, respectively) as seen in figure 2. We did detect type II methanotrophs (serine based), relatives of the *Verrucomicrobium Methylacidiphilium* [24] and *Proteobacteria Methylobacter tundripaludum* [25]. Relatives to the anaerobic methanotroph *Candidatus Methylomirabilis* oxyfera of the NC10 candidate phylum [26] were identified only in a few samples.

Coinciding with a depletion of oxygen, genome equivalence estimates indicate that in KT and LB genes for fermentation were very abundant and they increased in the bottom layers of the water columns, suggesting that organic substrates represented the main electron acceptors. Given the low contribution of tricarboxylic acid (TCA) cycle related genes and most electron-transport chain complexes including terminal oxidases, we infer a strictly anaerobic fermentation-based lifestyle in bottom layers of KT and LB. In all the samples, the key enzymes for the glycolysis (Embden-Meyerhof-Parnas) pathway were present in addition to the potential to convert pyruvate to acetyl–coenzyme A (acetyl-CoA). We identified that pyruvate-formate lyase (PFAM:02901) and pyruvate ferredoxin oxidoreductase (PFAM:01855) had a larger presence when compared to pyruvate dehydrogenase.

### Key findings from metagenome assembled genomes

To further validate the presence of metabolic potential such as methanotrophy and iron oxidation, genomes were reconstructed for the members of the under ice communities – an approach applied previously to numerous environments [27] [28] [29]. Contigs were binned into 462 bins with 57 bins passing our quality criteria of more than 60% completeness and less than 10% contamination as assessed by CheckM [30]. Each of these high-quality bins, from now on called metagenome assembled genomes (MAGs), represented a minor portion of the community in each sample as their average coverages ranged from 0.12 to 6.57% of the mapped reads. For more details, see the online reports in supplementary material. The energy metabolism of the lakes’ microbes was a combination of using all different electron donors available in the water column with the high quality bins representing a full range of microbial metabolisms from chemolithoautotrophy to photoheterotrophy as seen in figure 4.

**Figure 4.**
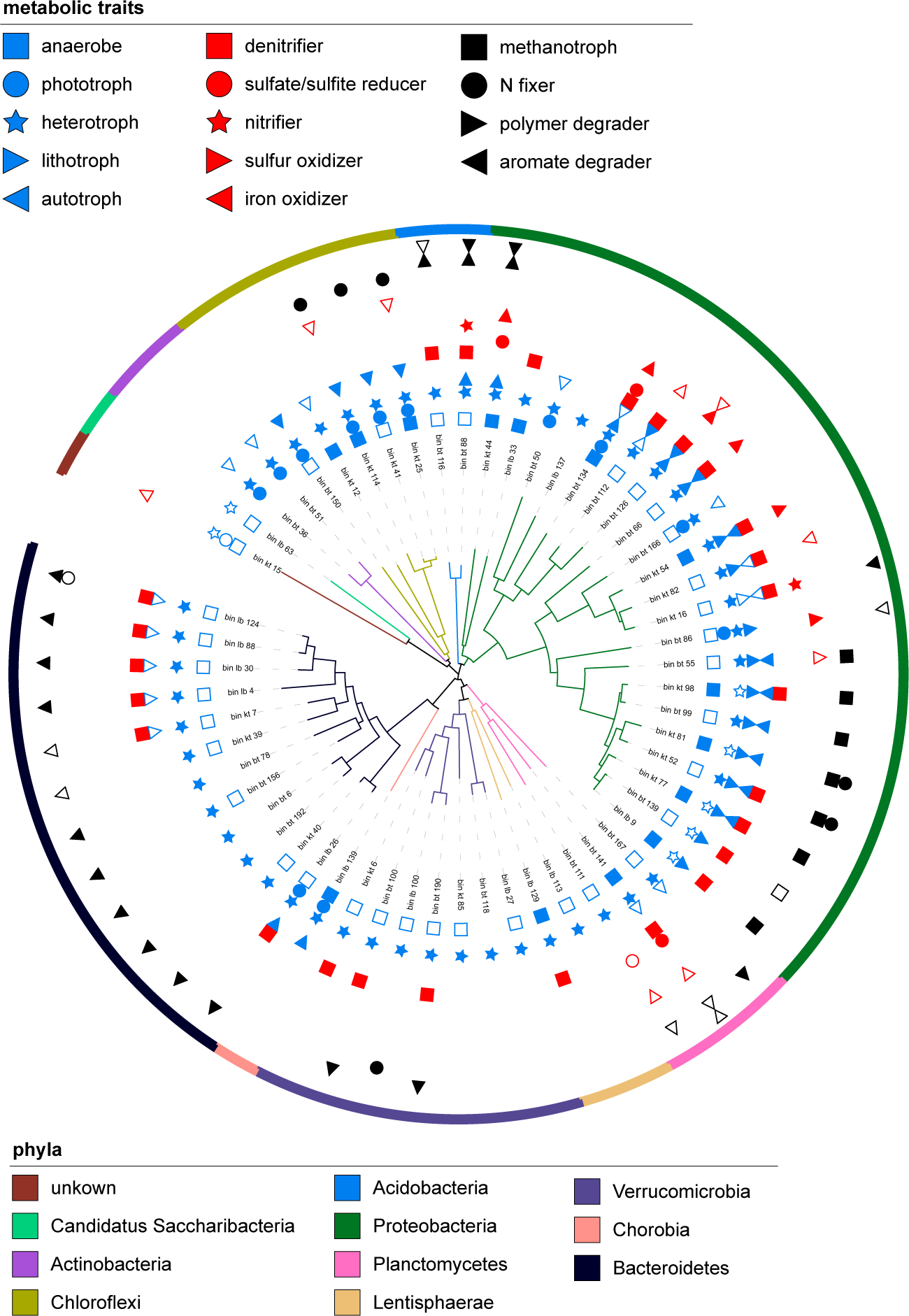
Traits, taxonomy and phylogeny of reconstructed genomes. Phylogenomic tree of recovered genomes as computed by PhyloPhlAn [79]. The outer ring onomic affiliation. The inner circles show the metabolic traits as based on genome raits with high support are indicated by full symbols, while traits with low support have open symbols. In the case a symbol is missing, no indications were found.

First, the 57 MAGs were taxonomically assigned to eleven phyla including Candidate phylum *Saccharibacter* (TM7), as seen in figure 4. When present in the MAG, the large and small subunit of the ribosomal RNA were assigned using the naive bayesian classifier [31] to obtain a more detailed taxonomic association, allowing the link with our amplicon data. After this assignment, the 57 MAGs were sent through multiple annotation pipelines to obtain a metabolic trait assignment, including their preferred electron acceptors and donors. Genomes of typical freshwater bacteria such as freshwater SAR11 (LD12) [32] and members of the acI clade [33] were obtained from the various systems with a high genome completeness. Annotations confirmed that, analogous to genomes from single cells and cultures [34] [35] [32], these contained the genes necessary for carrying out the Embden–Meyerhof–Parnas (EMP) pathway, the tricarboxylic acid (TCA) cycle, and a typical electron transport chain supplemented with light-mediated ATP production (i.e. bacteriothodopsin). Additional genomes including members of the phyla *Bacteriodetes*, *Acidobacteria* and *Plancomycetes*, contained a large variety of carbohydrate-active enzyme families, as shown in figure 4, indicating their involvement in allochthonous plant derived polymer degradation in the boreal lakes. For details, we refer the reader to the “MicroScope” pipeline [36] [37] with automatic annotations presented in KEGG maps and MetaCyc collections.

In addition to these heterotrophic degraders, we were able to obtain multiple MAGs related to previously described bacteria implicated in methanotrophy such as *Methylobacter tundripaludum*. The multiple genomes obtained related to *M. tundripaludum* contained *pmoABC* genes, and encoded for the complete reductive citrate cycle (Arnon-Buchanan cycle, rTCA) and *N*_2_-fixation [25] [38]. Since several genomes contained nitrate reductase (*nar*), nitrite reductase (*nir*) and nitric oxide reductase (*nor*) operons similar to genomes obtained from wetlands [25] [38], it can be speculated that these organisms combine denitrification with anaerobic methane oxidation, using a pathway similar to that of *Candidatus* Methylomirabilis oxyfera [26]. This pathway is proposed to include a quinol-dependent nitric oxide reductase (q*nor*) and a nitric oxide dismutase (*nod*) [39] with homologs identified in several of the *M. tundripaludum* MAGs. Even though it was shown that under hypoxia such methanotrophs can also use nitrate as terminal electron acceptor, oxygen seems to be necessary to activate methane by *pmo*, since no intrinsic oxygen production such as in *M. oxyfera* was observed [40]. Alternatively, the ability to grow heterotrophically was also implied by the presence of complete glycolysis and TCA cycle in the genomes. Other methanotrophs representing distinct lineages within the bacterial phylum *Verrucomicrobia*, such as *Candidatus Methylacidiphilum*, were present in the amplicon dataset but could not be retrieved in the genome reconstructions.

It is of note that a partial *Chlorobium* genome, with the closest relative identified to be *Chlorobium ferroxidans*, reconstructed from LB possessed neither a complete set of genes required for sulfur and iron oxidation nor all encoded components required for a photosynthetic electron transfer chain. In the iron-rich lake KT, the most representative bins were taxonomically assigned to the phylum *Chloroflexi*, most closely related to *Oscillochloris trichoides*. These bins encoded the complete reductive pentose phosphate cycle (Calvin-Benson-Bassham cycle, CBB). The fact that *Chloroflexi* can fix carbon dioxide by employing the CBB rather than by the 3-hydroxypropionate cycle was also recently shown on isolated *O. trichoides* producing a type I ribulose-1,5-bisphosphate carboxylase–oxygenase (RubisCO) [41]. In addition to a complete nitrogen fixing operon, several of the closely related Chloroflexi MAGs possessed a complete bacteriochlorophyll a and c synthesis pathways, allowing the prediction of phototrophic potential with an absorption maxima in the far red (750 and 860 nm) [42]. However, compared to previously described sulfite [41] and nitrite [43] oxidizing *Chloroflexi*, neither *sox*, *dsr* nor *nir* genes were present in our genomes. Like *O. trichoides* and *Chloroflexus aurantiacus* two Chloroflexi MAGs, ‘kt_25’ and ‘kt_114’, contained homologs to genes and gene clusters for the alternative complex III (ACIII), a cupredoxin (with redox potentials ranging from <200 mV (e.g aracyanin) to 700 mV in azurin [44]) and NADH:quinone oxidoreductase representing a third type of photosynthetic electron transport complex [45]. In addition homologs to a Rieske protein representing a widely ranging electron reduction potential from −150 to 400 mV, could be identified in ‘kt_25’ and ‘kt_41’. Considering that the redox potential of minerals such as FeS and FeCO3 is about +200 mV at pH 7, this together with the essential pieces resembling a ACIII reaction center with a membrane-spanning electron transfer chain terminating in Fe-S centres of the the Rieske protein rather than dissociable quinones could hypothetically generate reverse electron flow from Fe compounds (likely also Fe(II)) to NADH.

Under acidophilic conditions, few known microorganisms gain energy by the oxidation of Fe(II) and oxygen to generate reverse electron flow from Fe(II) to NADH [46]. Homologs to the redox cofactor pyrroloquinoline quinone, a protein with a transport function, iron oxidation specific c-type cytochrome *foxE* with no significant similarity to other known proteins and the cytochrome oxidase subunits I and II were identified in two representatives of the *Planctomycetes* phylum. In addition, multiple genomes with strong signals of chemolithoautotrophy could be derived from the three systems, including reduction of inorganic nitrogen and the oxidation of inorganic sulfur and nitrogen as seen in figure 4. Other interesting findings were genomes indicating high metabolic plasticity similar to *Rhodopseudomans palustris* as they seem to be capable of using multiple redox reactions within their population. These were *Betaproteobacteria* belonging to *Acetobacteraceae* and *Bradrhizobiaceae* with both chemolithoautotrophic and chemoorganoheterotrophic capability using oxygen, nitrate, Fe(III), sulfate and even organic compounds as electron acceptors. We also found an extremely coding-dense genome identified belonging to the *Candidatus Saccharibacteria* (TM7) phylum. This phylum has been suggested to contain epibionts or parasites of other bacteria due to their extreme auxotrophy [47].

## Discussion

Gross metabolic properties of the boreal lakes have been studied extensively indicating their netheterotrophy, as inferred from partial pressure of carbon dioxide [48] [49] [50]. If we want to predict and modulate ecosystem functions, knowledge of the quantities and types of organisms as well as their functions that constitute a particular ecosystem is an essential first step. With deep sequencing results presenting a detailed taxonomic inventory throughout the water column of five ice-covered boreal lakes, we show that the bacterial diversity differed widely among the lake systems. Taxonomic stratigraphy already shows some indications concerning the properties of the microbial communities inhabiting the redox towers, such as that anoxygenic phototrophs and methanotrophs are widely distributed in these ice-covered systems, similar to their icefree state [51]. Although sequences from well described branches of life, including certain methanotrophic and sulfur reducing Proteobacteria were observed, most of identified taxa are from less studied clades (e.g. representatives of phyla *Chloroflexi* and *Planctomycetes*). These clades populate a large part of phylogenetic tree of life, yet are all in poorly documented areas where we have very limited knowledge about the metabolic repertoire found in their genomes.

By exploring microbial genomes in a high throughput fashion we show that a wide range of metabolic capabilities involving the use of multiple electron donors and acceptors appears to be common in the microbial community in the studied lake systems. In spite of redox metabolic plasticity occurring in microbial community of each system, we found that the majority of organisms probably lack the ability to perform multiple sequential redox transformations within a pathway. Further, we revealed variations at the genomic level within and between systems when it came to iron and sulfur cycling, which is further emphasized by specific taxonomic groups proliferating in each lake system. Our genomic data exposes diverse metabolic traits, such as photoferrotrophy, chemolithotrophy and methane oxidation, supporting alternative energy sources in these net-heterotrophic ecosystems, as seen in figure 5. From this first detailed genetic characterization, a complex metabolic system emerges that can be hypothesized to depend on the availability of light as determined by organic matter load, ice thickness, snow cover and weather conditions, as well as the availability of nutrients (e.g. N, P, S, Fe) as determined by the catchment, and, last but not least, by the physical properties (i.e. morphology and hydrology).

**Figure 5.**
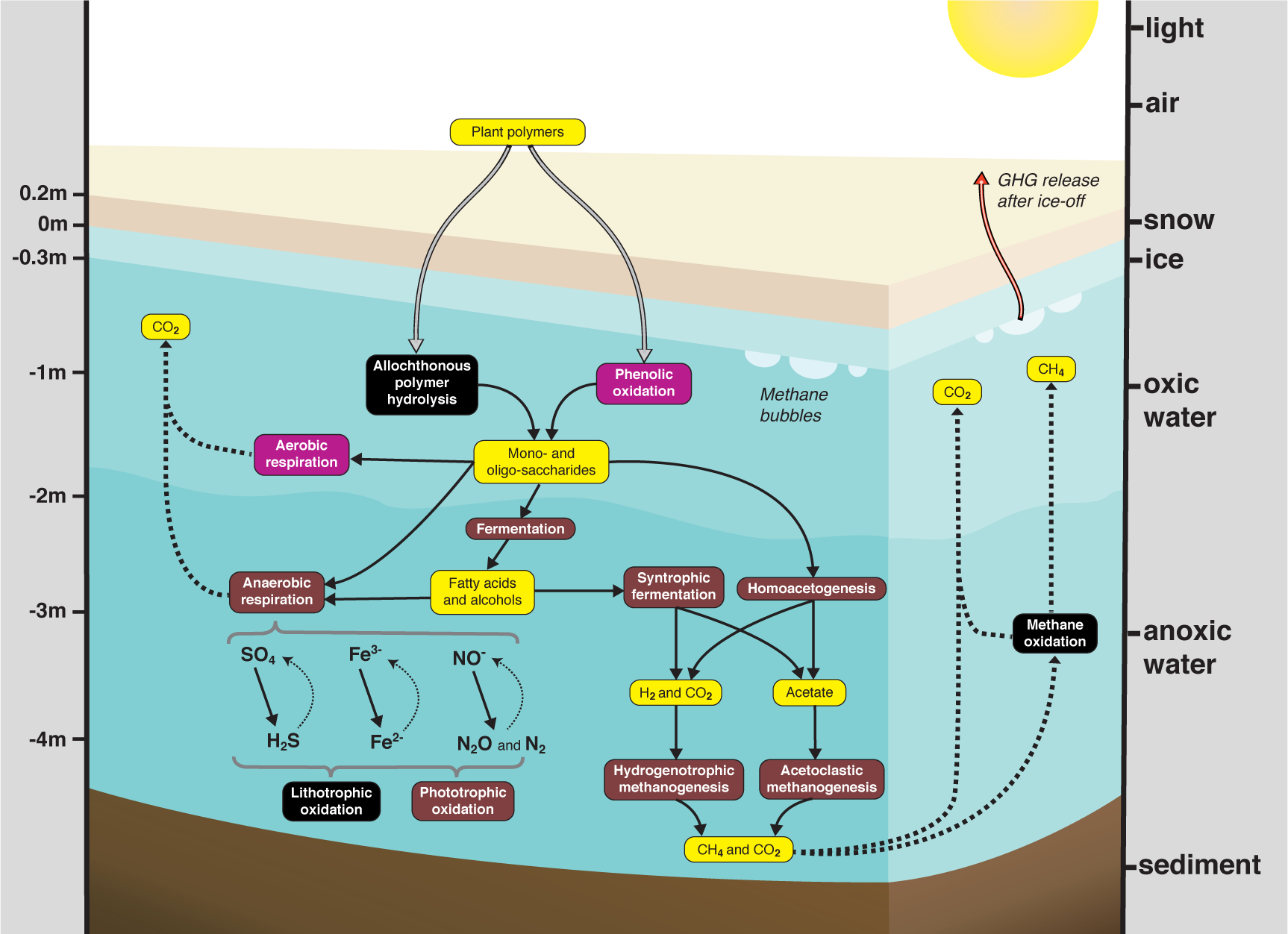
Main degradation pathways of allochthonous organic matter. Schematic overview of the main degradation pathways of allochthonous organic matter occurring under ice in boreal lakes. The pathways are divided into three categories: aerobic (purple), anaerobic (brown) and processes occurring under both conditions (black). Compounds are represented in yellow. Metagenome assembled genomes (bins) encoding for the different metabolic processes are presented in figure 4. The figure is adapted from figure 1 of Conrad and colleagues [80].

As we predict that organisms mediate individual reaction steps in redox pathways these must be linked to form full or short-circuit cycles of elements. Short circuits are for example elemental cycles where the most reduced and oxidized forms are not the most common reaction products. Instead, interconversions of sulfide to elemental sulfur or denitrification from nitrate to nitrite, resulting in products that can be oxidized back to sulfide and nitrate by phototrophs or chemilithotrophs are very commonly encoded in the genomes. Such restricted metabolic potential may give these highly specialized organisms an advantage under the redox conditions at certain depth, even though organisms related to for example *Rhodopseudomans palustris* with a wide range of metabolic capabilities do co-exist. Overall, the reconstructed depth profile of the lakes allowed us to estimate the degree with which specialized metabolic niches are formed and to quantify the proportion of genomes with the potential to obtain their energy from either light, oxidation of organic molecules, or the oxidation of inorganic molecules.

Our findings emphasize that elemental cycles, such as those of sulfur and iron, have the potential to play major roles in the systems and need to be taken into consideration when building biogeochemical models. By contrast, nitrogen cycling seemed to be less widely distributed among the microbes inhabiting frozen boreal lakes, as previously thought, while the importance of methanotrophy-driven systems is corroborated [52]. Based on stable isotope experiments, a pelagic food web largely supported by methane metabolism was already proposed previously [52]. Photoferrotrophy has been shown to play a role in a handful of systems only very recently [53] [54] [55]. These systems have been regarded as modern analogues of the Archaean ocean in a time where oxygenic phototrophs had yet not evolved, as both photoferrotrophy and anaerobic methane cycling are present. Our analyses based on genomic reconstructions in conjunction with a very recent publication using isotope approaches [11] suggest that anoxic iron and methane cycling may be widely distributed among the millions of lakes in the boreal zone.

To sum up, the insight gained from the reconstruction of the microbial genomes lead us to formulate hypotheses concerning the biogeochemical cycling of elements in these ecosystems of global significance. We provide an overview of the ongoing metabolic processes as shown in figure 5, which resembles most redox stratified systems. Besides some general resemblance, metabolic and taxonomic profiles emerged unique to each lake’s redox tower and with sulfur, iron and carbon cycling tightly intertwined through chemolithotrophy and anoxygenic photosynthesis.

## Materials and Methods

### Sample design and environmental properties

In total, thirty-nine water samples were obtained from five boreal lakes (seven or eight samples per lake) all located in central Sweden. Our sampling took place in the area covered by N63°33’ to N63°59’ and E12°27’ to E14°46’ during March 18th to 20th of 2014. Each lake was assigned a two letter acronym for easy reference: RL, BT, KT (Lomtjärnen), LB (Liltjärnen) and SB (Fröåtjärnen) as some of the lakes posses no authoritative names to our knowledge. These water samples were obtained along a depth gradient by drilling small holes on the ice surface and by using a rope-operated Limnos sampler. Samples were immediately processed: for *CO*_2_ and *CH*_4_, 25 ml of samples were directly taken from the sampler with 60 ml polypropylene syringes equipped with three-way stopcocks and subsequently stored on ice. Upon return to the laboratory, after creating a 10 ml room-air headspace in each syringe, *CH*_4_ was equilibrated between the remaining water and the headspace by vigorous shaking for 3 minutes. The headspace was then injected into *N*_2_ filled 120 ml infusion vials with crimp aluminium seal secured 10-mm butylrubber stoppers [56]. Within a week, gas concentrations were measured with a gas chromatograph (GC) (Agilent Technologies 7890A GC Systems) equipped with a flame ionization detector by injecting 1 ml of gas from the infusion vials. *CH*_4_ concentrations were calculated according to Henry’s law, correcting for temperature according to [57] and corrected for *CH*_4_ in ambient air as measured on the GC.

Water temperature and oxygen concentrations were measured *in situ* using a YSI 55 combined temperature and oxygen probe (Yellow Springs Instruments, Yellow Springs, Ohio, USA). Total phosphorus (TP) and total nitrogen (TN) were measured using standard methods as previously described [58]. Total organic carbon (TOC) concentrations, also known as non-purgeable organic carbon (NPOC), were obtained by analysis on a Shimadzu TOC-L with sample changer ASI-L (Shimadzu Corporation, Japan).

Water for analyses of 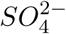 analysis was first pre-filtered through rinsed 0.2 µm membrane filters (Pall Corporation) and then analyzed by ion chromatography on a Metrohm IC system (883 Basic IC plus and 919 Autosampler Plus) fitted with a Metrosep A Supp 4/5 guard column and a Metrosep A Supp 5 analytical column (150x4.0 mm). The concentrations of Fe(II) and Fe(III) were measured with the ferrozine colorimetric method [59].

Samples for bacterial counts were fixed with 37% borax buffered formaldehyde (final concentration 2%) and stored at 4 °C prior to analyses. Cells were stained with the fluorescent nucleic acid stain Syto13 (Molecular probes, Invitrogen, Carlsbad, CA, USA) according to the protocol of Del Giorgio and colleagues [60] and were then counted with a flow cytometer equipped with a 488 nm blue solid state laser (Cyflow Space, Partec, Görlitz, Germany) using green fluorescence for triggered particle scoring. Cell counts were analyzed using Flowing Software version 2.5 (Perttu Terho, Centre for Biotechnology, Turku, Finland).

### DNA extraction and 16S rRNA gene amplicon sequencing

For DNA, between 0.5 and 1 liter of water was filtered through 0.2 μm Sterivex cartridges (Millipore), in duplicate, by using sterile syringes followed by on site liquid nitrogen freezing until further analyses. After filters were removed from the cartridges, total DNA was extracted using a Powersoil DNA Isolation Kit (MO BIO). Bacterial 16S rRNA gene amplicons were sequenced on a MiSeq machine (Illumina) following procedures modified from Sinclair and colleagues [61]. In short, each sample was first amplified in duplicate using primers targeting the variable regions of the rRNA gene (V3/V4 region) and equipped with parts of the Thruplex Illumina sequencing adapter. After duplicates were pooled and purified using the Agencourt AMPure XP system (Beckman Coulter) as recommended by the manufacturer, the pooled samples were used as templates in a second PCR step using primers equipped with a 7bp index and the Illumina sequencing adapters for multiplexing. After purifying the samples using the Agencourt AM-Pure XP kit and quantifying by fluorescence with the PicoGreen assay (Quant-iT PicoGReen, Invitrogen), samples were pooled in equimolar amounts. The pooled samples were sequenced at the SciLifeLab SNP/SEQ sequencing facility (Uppsala University, Uppsala, Sweden) using an Illumina MiSeq with a 2x300 bp chemistry. Finally, a 16S analysis and annotation pipeline, previously described in [61], was used. The sequence processing comprises steps for pairing reads, quality filtering, chimera checking, clustering with a 3% sequence dissimilarity threshold, taxonomic assignment, diversity estimations and result visualization.

### Shotgun-metagenomic sequencing

First, 10 ng of genomic DNA was sheared using a focused-ultrasonicator (Covaris E220). Next, the sequencing libraries were prepared with the Thruplex FD Prep kit from Rubicon Genomics according to the manufacture’s protocol (R40048-08, QAM-094-002). The library size selection was carried out with AMPure XP beads (Beckman Coulter) in a 1:1 ratio. The prepared sample libraries were quantified by applying KAPA Biosystem’s next-generation sequencing library qPCR kit and run on a StepOnePlus (Life Technologies) real-time PCR instrument. The quantified libraries were then prepared for sequencing on the Illumina HiSeq sequencing platform utilizing a TruSeq paired-end cluster kit, v3, and Illumina’s cBot instrument to generate a clustered flowcell for sequencing. Two individual sequencing runs (an evaluation and re-run) were performed with twenty-four samples. In both cases, sequencing of the flowcell was performed on the Illumina HiSeq2500 sequencer using Illumina TruSeq SBS sequencing kits, v3, with the exception of following a 2×100 bp indexed high-output run recipe for the evaluation and a 2×125 bp in the case of the re-run.

### Shotgun-metagenome processing

Reads were filtered based on their PHRED quality scores using sickle (version 1.33) [62] and then assembled with Ray (version 2.3.1) [63]. Prior to generating a final assembly, we created a number of assemblies for optimization on the evaluation run. Assemblies with different k-mer sizes (31-81) were compared, based on different metrics such as N50. Assemblies of k-mer sizes of 51, 61, 71 and 81 were chosen to be applied on the re-run. Our tests further evidenced that when contigs were pooled and cut into 1’000 bp pieces and reassembled with Newbler (version 2.9) (454 Life Sciences, Roche Diagnostics) N50 was increased, therefore we used this technique to produce the final assembly on which all analyses were performed. Coverage was computed by running bowtie (version 2.2.5) [64] to map the reads back to the Newbler-produced assembly. Duplicates were removed by picard-tools (version 1.101). For computing coverage, bedtools (version 2.18.2) [65] was used. Once contigs and scaffolds shorter than 1 kb were discarded, concoct (version 0.3.0) [66] was run for binning. Bins were separated into low quality and high quality groups, based on results obtained from CheckM (version 0.9.7) [30]. A completeness value of over 60% and a contamination metric below 10% were the criteria chosen for classifying a bin as ‘good’ (high quality) and for it to obtain the denomination of “metagenome assembled genome” (MAG).

### Functional trait profiles

Hmmsearch (version 3.1b2) [67] on the PFAM-A database at version 29.0 [68] was used to provide the annotation of the proteins predicted by Prodigal (version 2.6.2). Coverage information and abundance of proteins allowed us to estimate genome equivalence of individual PFAMs by normalizing each individual PFAM with the average quantity of 139 PFAMs predicted to occur in single copy in all genomes [69].

### Genome analysis

Genome annotations were performed using “MicroScope” with automatic annotations assisted by manual curation, as described in the integrated bioinformatics tools and the proposed annotation rules [36] [37]. In addition to the integrated annotation tools, which includes BlastP homology searches against the full non-redundant protein sequence databank, UniProt [70] and against the well-annotated 164 model organisms *Escherichia coli K-12* and *Bacillus subtilis* 168 [37], enzymatic classifications based on COG [71], InterPro [72], FIGFam [73] and PRIAM [74] profiles, and prediction of protein localization using the TMHMM [75], SignalP [76] and PSORTb [77] tools, we also used our local hmmsearch results. Synteny maps (i.e. conservation of local gene order) were used to validate the annotation of genes located within conserved operons. Metabolic pathways were subsequently identified with the assistance of the integrated MicroCyc database and the Kyoto Encyclopedia of Genes and Genomes (KEGG) database [78]. Genomes can be retrieved from “MicroScope” with their specific identifiers as given in the supplementary material.

### Statistical analyses

Statistical analyses were done using Python packages or the R language, including non-parametric multidimensional scaling (NMDS) plots, permuted multivariate ANOVA (PERMANOVA), redundancy analysis and the Procrustes superimpositions. The analyses were performed on rarefied OTU tables of amplicon data using the Bray-Curtis distance and PFAM tables standardized to genome equivalents [69] using the Morisita-Horn distance. Particular attention was paid to pathways linked to energy metabolism and carbon cycle when analyzing metabolic depth profiles. To plot the depth profiles of the traits, marker HMMs were selected as unique members of specific traits and are listed in supplementary table S1.

## Acknowledgments

We would like to thank the various institutions running the multiple computational resources that were used in this project: (i) the Swedish National Infrastructure for Computing (SNIC) through the Uppsala Multidisciplinary Center for Advanced Computational Science (UPPMAX), (ii) the IT Center for Science in Finland (CSC) and (iii) the Cloud Infrastructure for Microbial Bioinformatics (CLIMB) funded by the UK’s Medical Research Council (MRC).

We also would like to acknowledge the support given by the SciLifeLab SNP/SEQ facility hosted by Uppsala University in the molecular sequencing steps.

This research was made possible through a scholarship from the Olsson-Borgh foundation for limnological studies (Uppsala University foundation number 91173 to LS) as well as grants by the Swedish Research council (grant 2012-4592 to AE), the Swedish Foundation for strategic research (grant ICA10-0015 to AE) and the Academy of Finland (grant 265902 to SP).

This research was carried out at Uppsala University in the Department of Ecology and Genetics, Limnology and manipulations were done in the laboratory of the Evolution Biology Center at Norbyvägen 18E, Uppsala, Sweden.

We would like to thank Moritz Buck for his input on this manuscript and the bioinformatics methods used.

## Author contributions

All authors participated in revising the manuscript.

**LS**: Designed the study. Participated in sampling. Processed the sequence data and did the bioinformatics. Carried out statistical analyses. Wrote this manuscript.

**SP**: Conducted and orchestrated the sampling campaign. Processed samples in the lab.

**MB**: Participated in sampling. Provided advice on bioinformatics procedures and methods.

**PH**: Participated in sampling. Processed samples in the lab.

**MS**: Participated in sampling.

**AE**: Designed the study. Participated in sampling. Processed the sequence data. Carried out statistical analyses. Wrote this manuscript.

The authors declare no competing financial interests or other conflicts of interest.

## Supporting Information

**Table S1.** Selected PFAMs traits.

Supplementary table S1 is too long and not shown here. It is suited for online distribution after publication and to be viewed within a spreadsheet application. It details the list of PFAMs chosen to be unique to specific pathways.

**Table S2.** PFAMs found per bin.

Supplementary table S2 is too large to be shown here. It is best suited for online distribution after publication and to be viewed with a spreadsheet application. It shows the results of the hmmsearch against the PFAM database for lakes BT, KT and LB. Columns are bin identifier numbers and rows are PFAMS. Each value indicates how many times each PFAM was found in each bin.

**Figure S1.**
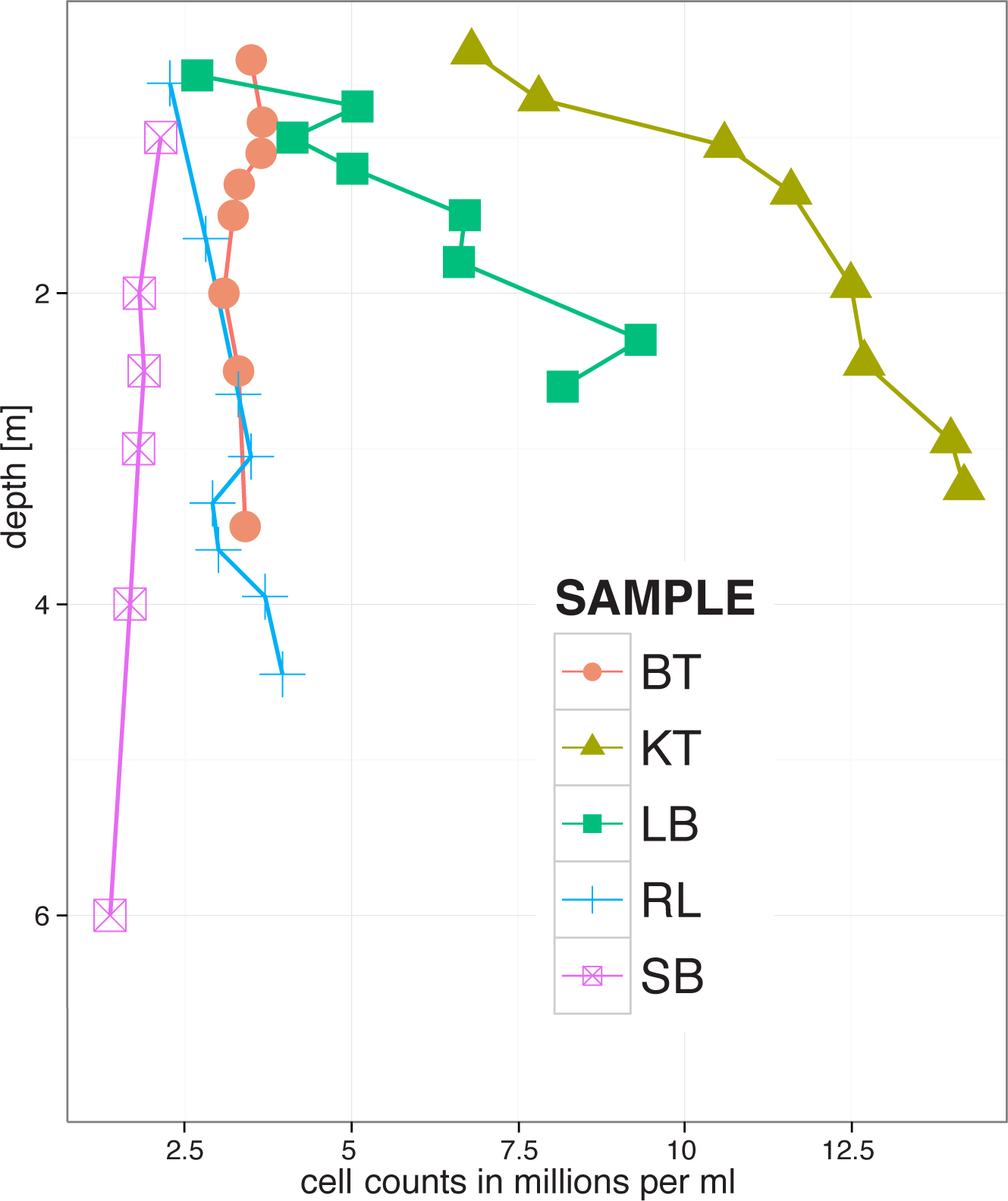
Cell counts per depth for all lakes.

**Figure S2.**
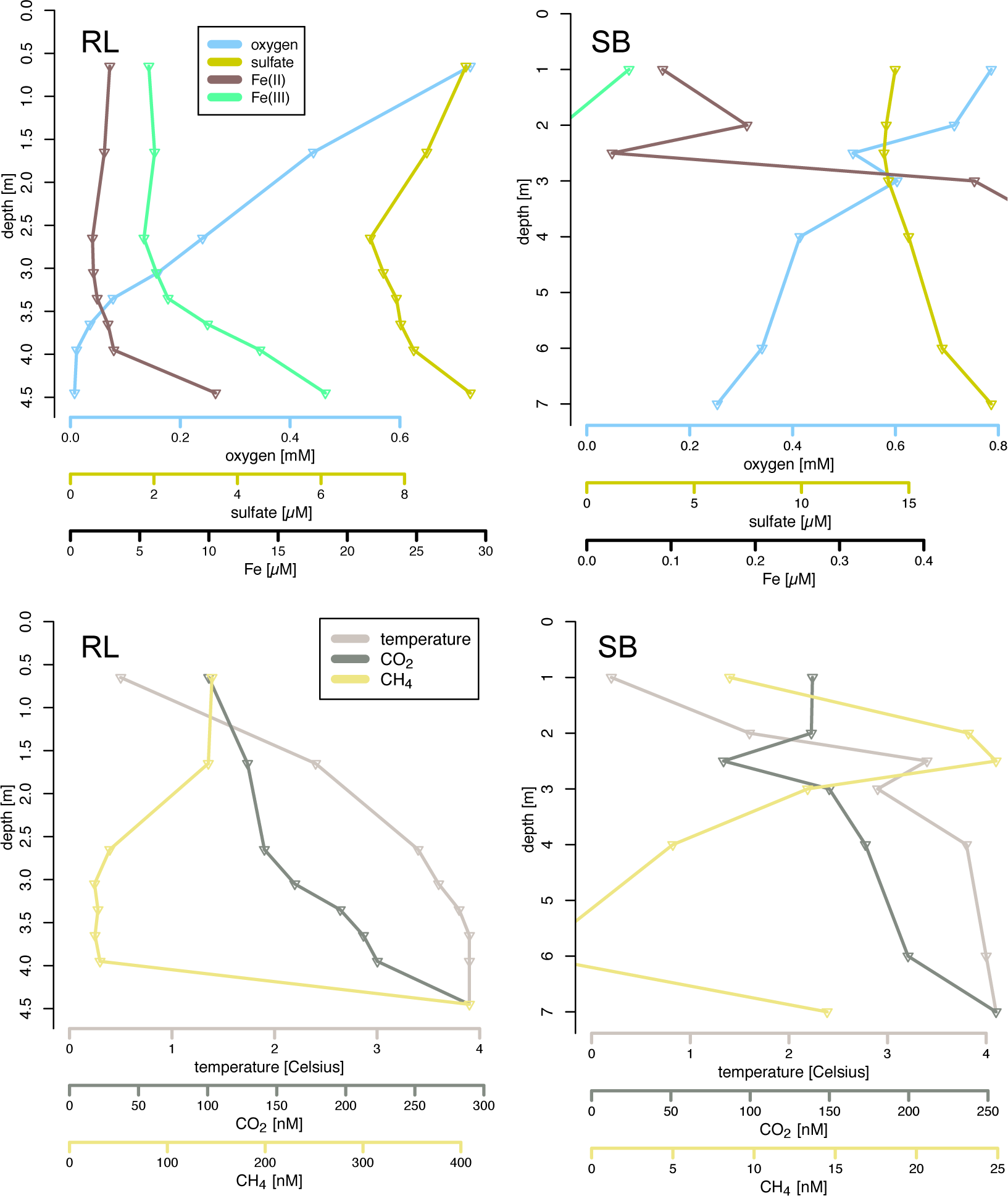
Chemistry and gas profiles in lakes RL and SB.

**Figure S3.**
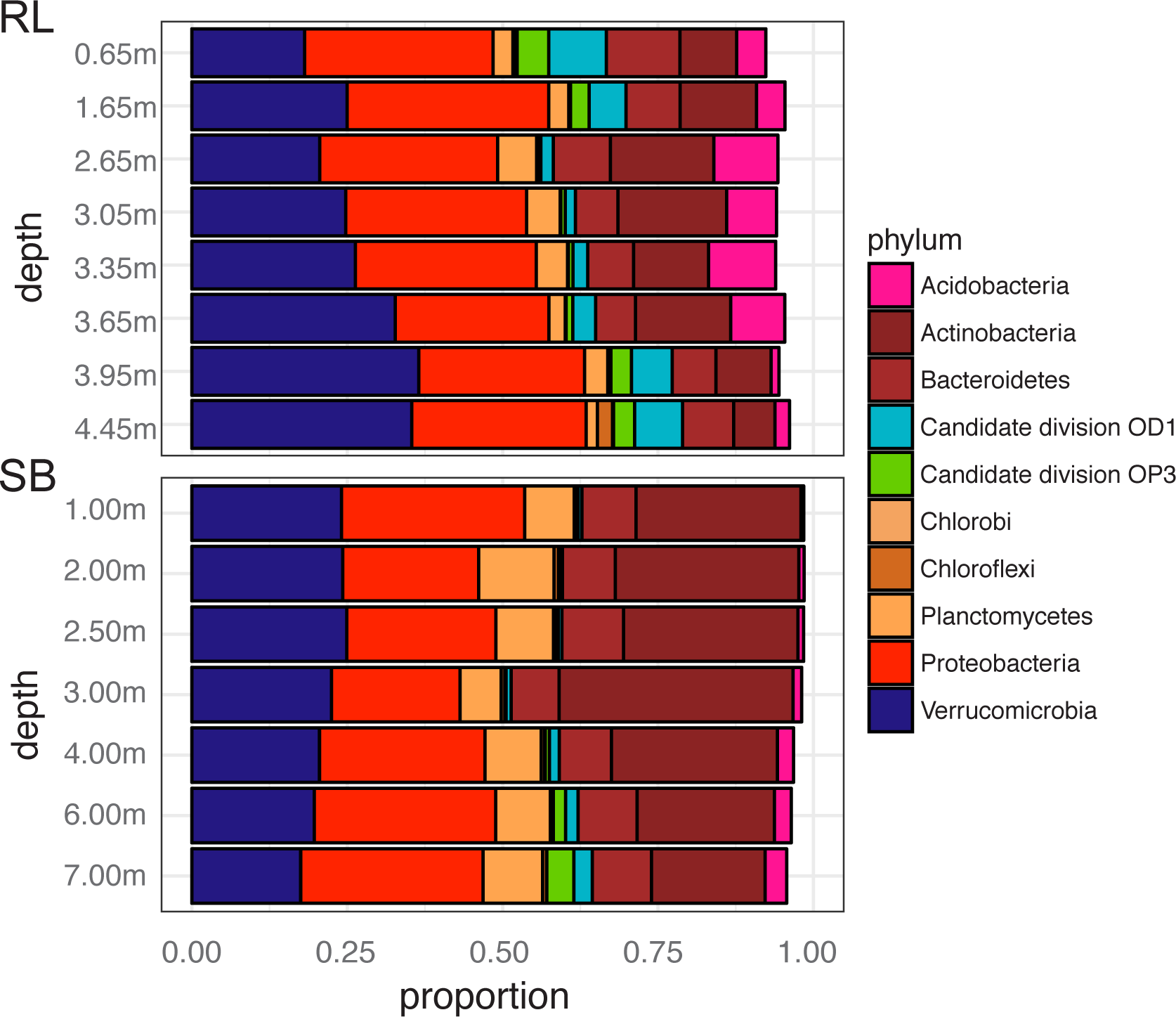
The distribution of the ten most abundant phyla in lakes RL and SB.

**Figure S4.**
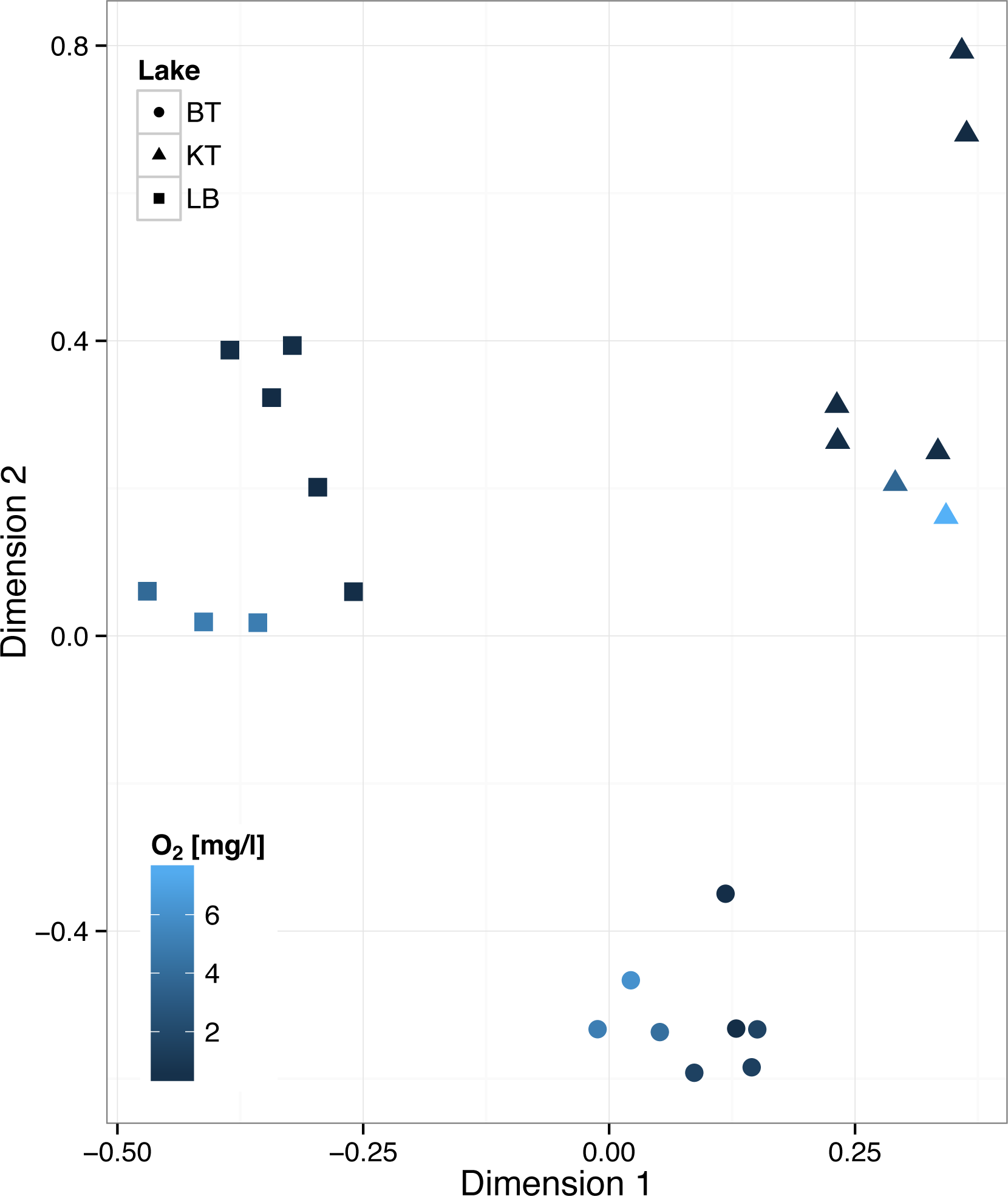
NMDS based on PFAMs found in three lakes.

